# Clinical isolates of *Mycobacterium avium* complex reveal an *M. intracellulare*-associated IL-17/neutrophilic pulmonary immune program in a murine disease model

**DOI:** 10.64898/2026.05.27.728352

**Authors:** Shintaro Seto, Koji Furuuchi, Shiho Omori, Hajime Nakamura, Kozo Morimoto, Minako Hijikata, Naoto Keicho

**Author notes:** Correspondence: Shintaro Seto.

## Abstract

*Mycobacterium avium* complex (MAC) is the leading cause of nontuberculous mycobacterial pulmonary disease (NTM-PD) and mainly comprises *M. avium* (MAV) and *M. intracellulare* (MI). Host–pathogen interactions may contribute to the heterogeneous clinical course of MAC pulmonary disease (MAC-PD); however, species- or isolate-associated differences in virulence and host immune responses induced by MAC strains remain poorly understood. Here, we established a panel of MAC clinical isolates exhibiting persistent pulmonary infection in mice and performed transcriptomic analyses to evaluate pulmonary immune responses. Although MAC infections induced broadly shared inflammatory responses, MI infection elicited a robust IL-17/neutrophilic inflammatory signature, whereas MAV infection showed relative enrichment of IFN-γ/cytotoxicity-associated responses, indicating differences in the balance of their immune gene expression programs. The MI-associated IL-17/neutrophilic program remained evident at the isolate level and after adjustment for pulmonary growth phenotype. RT-qPCR and flow cytometric analyses using the representative isolate pair FKJ-1 (MI) and FKJ-8 (MAV) further supported the differential induction of these immune signatures in infected lungs. To define the cellular basis of these immune programs, we performed single-cell RNA sequencing on lungs infected with FKJ-1 and FKJ-8. Infection with the representative MI isolate was associated with increased *Il17a*-expressing CD4^+^ T cells and γδ T cells, neutrophilic inflammation, and the expansion of inflammatory macrophages. Taken together, these findings demonstrate that within a selected panel of persistent MAC clinical isolates, MI infection is associated with a robust IL-17/neutrophilic pulmonary immune program, providing a preclinical framework for dissecting species- and isolate-associated host–pathogen interactions in MAC-PD.

## INTRODUCTION

Nontuberculous mycobacteria (NTM) are environmental mycobacteria increasingly recognized as an important cause of chronic pulmonary infections (1). The burden of NTM pulmonary disease (NTM-PD) has risen in several regions, including Japan (2–4). Although the distribution of causative NTM species varies by geographic region, the *Mycobacterium avium* complex (MAC) remains the predominant cause of NTM-PD in Japan and globally (1, 2). NTM-PD, including MAC pulmonary disease (MAC-PD), is associated with diverse host factors, such as structural lung abnormalities and impaired host defenses; however, it is also frequently observed in patients without overt immunodeficiency (5, 6). These observations suggest that disease development is shaped by complex interactions among host susceptibility, the airway microenvironment, bacterial factors, and environmental exposures (7, 8).

NTM-PD, including MAC-PD, presents with highly heterogeneous clinical and radiological features. Radiologically, NTM-PD is commonly classified into fibrocavitary (FC) and nodular bronchiectatic (NB) types (1, 9). The FC type, characterized by cavitary and destructive lung lesions, is associated with poor treatment responses and prognosis. In contrast, the NB type is characterized by bronchiectasis, nodular lesions, and a relatively indolent disease progression. However, these phenotypes are not mutually exclusive; several patients exhibit overlapping features, and some progress from NB-type disease to cavitary disease. Thus, these radiological forms represent overlapping phenotypes within a broader spectrum of chronic pulmonary disease.

Host immune responses play a central role in determining the progression of MAC-PD. Previous studies have implicated inflammatory cytokine responses, interferon (IFN)-associated pathways, Th1/Th17 immune axes, neutrophilic inflammation, and macrophage activation in MAC infection and subsequent disease progression (10–14). In human MAC-PD, transcriptomic analyses have also suggested that peripheral blood immune signatures can reflect disease activity or severity (15). Nevertheless, the association between different MAC species or clinical isolates and pulmonary immune programs remains unclear.

Clinical studies have reported possible differences in disease manifestations and outcomes between *M. avium* (MAV)- and *M. intracellulare* (MI)-associated MAC-PD, although the clinical impact of species differentiation may vary across cohorts (16–18). This inconsistency likely reflects the overall complexity of MAC-PD, in which host susceptibility, structural lung disease, environmental exposure, treatment history, and bacterial strain diversity together influence clinical progression. Therefore, whether MAV and MI intrinsically differ in the pulmonary immune states induced by them remains unclear. A controlled murine infection model using multiple persistent MAV and MI isolates provides a useful framework to address this question by comparing host immune responses under standardized experimental conditions.

Murine models have been widely utilized to study chronic MAC infection and treatment responses *in vivo*. These models have demonstrated that MAC isolates differ in their ability to persist, proliferate, and induce lung pathology, including granulomatous inflammation and necrotizing lesions (10, 13, 19–24). However, immunological mechanisms underlying strain- or species-associated disease heterogeneity remain unclear.

In this study, we established and characterized a panel of MAC clinical isolates that can induce persistent pulmonary infection in BALB/c mice. To compare species-associated pulmonary immune programs using this panel, we performed bulk RNA sequencing (RNA-seq) of lungs infected with multiple MAV and MI isolates under controlled experimental conditions. To validate these immune programs, we further performed RT-qPCR and flow cytometry using the representative isolate pair FKJ-1 (MI) and FKJ-8 (MAV) and defined their cellular basis using single-cell RNA-seq (scRNA-seq). Broadly shared inflammatory responses induced by MAC infection were identified, together with species-associated differences in the relative dominance of specific immune modules. This study provides a preclinical framework for dissecting MAC species- and strain-associated host responses, supporting future studies into therapeutic and diagnostic strategies for MAC-PD.

## RESULTS

### Identification of MAC isolates exhibiting persistent pulmonary infection in a murine MAC-PD model

We developed a screening system to identify MAC isolates with persistent pulmonary bacterial burdens in infected mice by combining genome-guided pooled infection with whole-genome sequencing (WGS), as described previously (19). Among the 39 clinical MAC isolates tested, three isolates, FKJ-3.1, FKJ-14, and FKJ-27, were identified as persistent candidates in infected mouse lungs. WGS-based phylogenetic analysis revealed that these newly identified isolates are genetically distinct from the strains previously used to establish murine MAC-PD models (19, 20) (Fig. 1A). Individual infection experiments confirmed that these isolates exhibited persistent or proliferative phenotypes in the murine pulmonary infection model (Fig. 1B and Fig. S1). Histological analysis further demonstrated the formation of peribronchial granulomatous lesions in the infected lungs (Fig. 1C). Taken together, these results expand the panel of MAC isolates capable of establishing persistent pulmonary infections in BALB/c mice, enabling subsequent comparisons of host responses induced by MAV and MI isolates. Importantly, this panel was selected on pulmonary persistence in mice and was not intended to represent the full clinical or genetic diversity of MAV and MI isolates.

**FIG 1.**
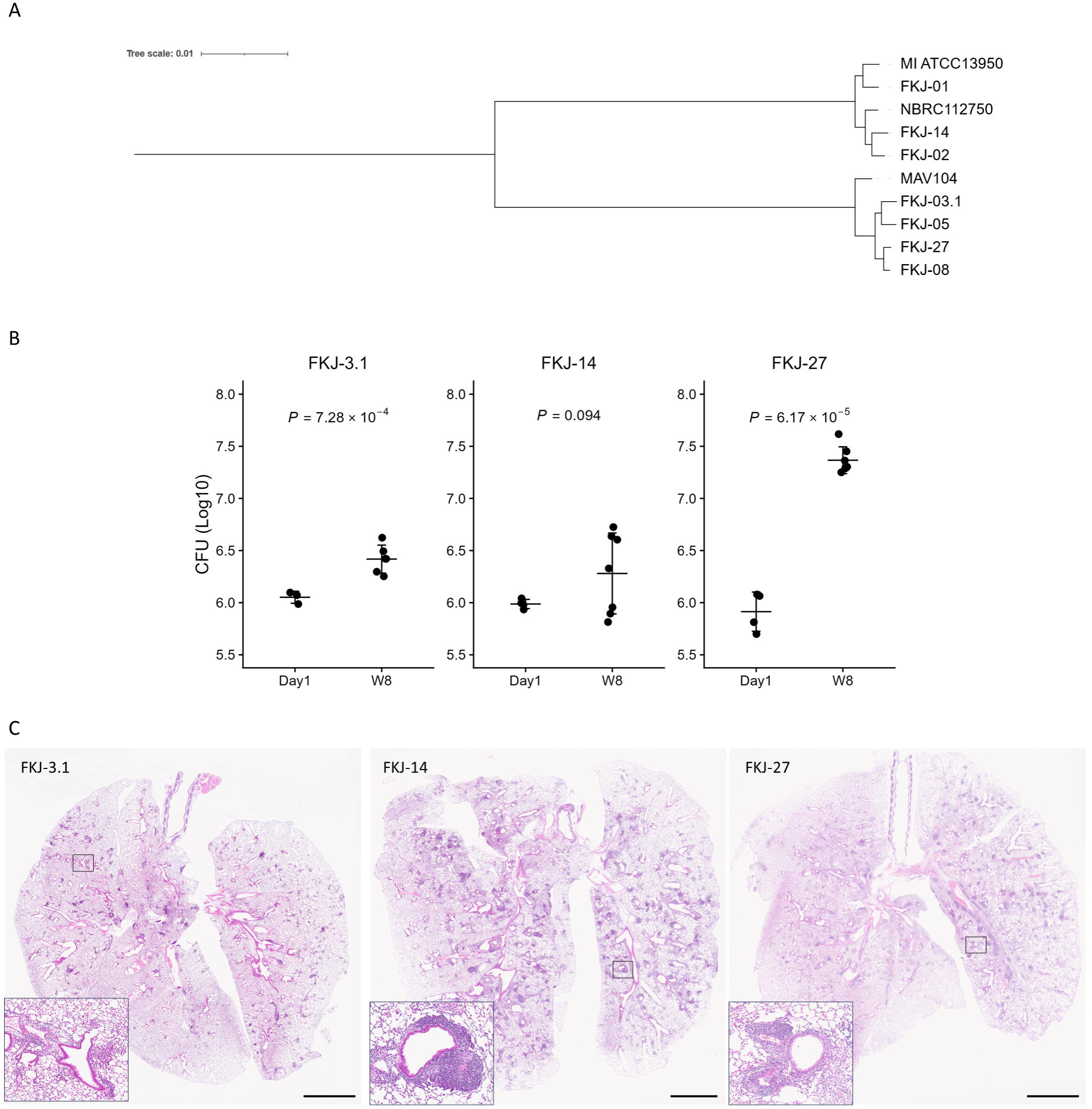
*Mycobacterium avium* complex (MAC) clinical isolates exhibiting persistence in a murine pulmonary infection model. (A) Phylogenetic tree of the MAC isolates used in this study. MAV104, *M. Avium* 104; MI, *M. intracellulare*. (B) Lung bacterial burdens in BALB/c mice infected with the clinical MAC isolates FKJ-3.1, FKJ-14, and FKJ-27. Bacterial burdens were examined at day 1 (Day1, n = 4) and 8 weeks (8W, n = 6) postinfection (p.i.). Each point represents an individual mouse, and horizontal bars indicate the mean. *P* values were determined using Welch’s *t*-test. (C) Representative hematoxylin and eosin-stained images of BALB/c mouse lungs infected with the indicated MAC isolates at 8 weeks p.i. Boxed regions are enlarged in the insets. Scale bars, 2.5 mm.

### Bulk RNA-seq reveals infection-induced differentially expressed genes in MAC-infected lungs

To evaluate pulmonary immune responses to MAC infection, we examined transcriptomic profiles in a chronic murine MAC-PD model. BALB/c mice were individually infected for 8 weeks with a panel of eight MAC isolates, comprising four MAV isolates, FKJ-3.1, FKJ-5, FKJ-8, and FKJ-27, and four MI isolates, FKJ-1, FKJ-2, FKJ-14, and NBRC112750. Bulk RNA-seq was then performed on the infected lung tissue. Compared with uninfected control lungs, MAC-infected lungs exhibited significant upregulation of genes associated with inflammatory responses, including type I and II IFN responses and TNF-α and IL-6 signaling pathways (Fig. S2A), consistent with previous reports (10, 13, 14). These infection-induced transcriptional changes were broadly observed in both MAV- and MI-infected lungs when each group was compared with uninfected controls (Fig. S2B). These results indicate that MAC infection induces common inflammatory and IFN-related gene expression programs in the lungs of this murine MAC-PD model.

Next, to identify species-associated differences in pulmonary immune responses within the eight-isolate panel, we compared the gene expression profiles of MAV-infected lungs with those of MI-infected lungs. Subsequently, 716 and 916 differentially expressed genes (DEGs) were found to be upregulated in MAV- and MI-infected lungs, respectively (Fig. 2A and Table S1). Gene set enrichment analysis (GSEA) revealed an enrichment of IFN-related pathways in MAV-infected lungs, whereas TNF-α and IL-6 signaling pathways were enriched in MI-infected lungs (Fig. 2B). Metabolism-associated Hallmark pathways, including mTORC1 signaling, hypoxia, cholesterol homeostasis, and glycolysis, were also enriched in MI-infected lungs compared with MAV-infected lungs. Gene Ontology biological process (GOBP) enrichment analysis of the upregulated DEGs further supported the presence of distinct transcriptional programs between MAV and MI infection (Fig. 2C). Genes associated with type I and type II IFN responses were enriched in MAV-infected lungs, whereas genes associated with IL-17 responses and neutrophil chemotaxis were enriched in MI-infected lungs. These results suggest that, within the isolate panel used in this study, MAV and MI infections are associated with distinct transcriptional programs in the lungs of this chronic murine MAC-PD model.

**FIG 2.**
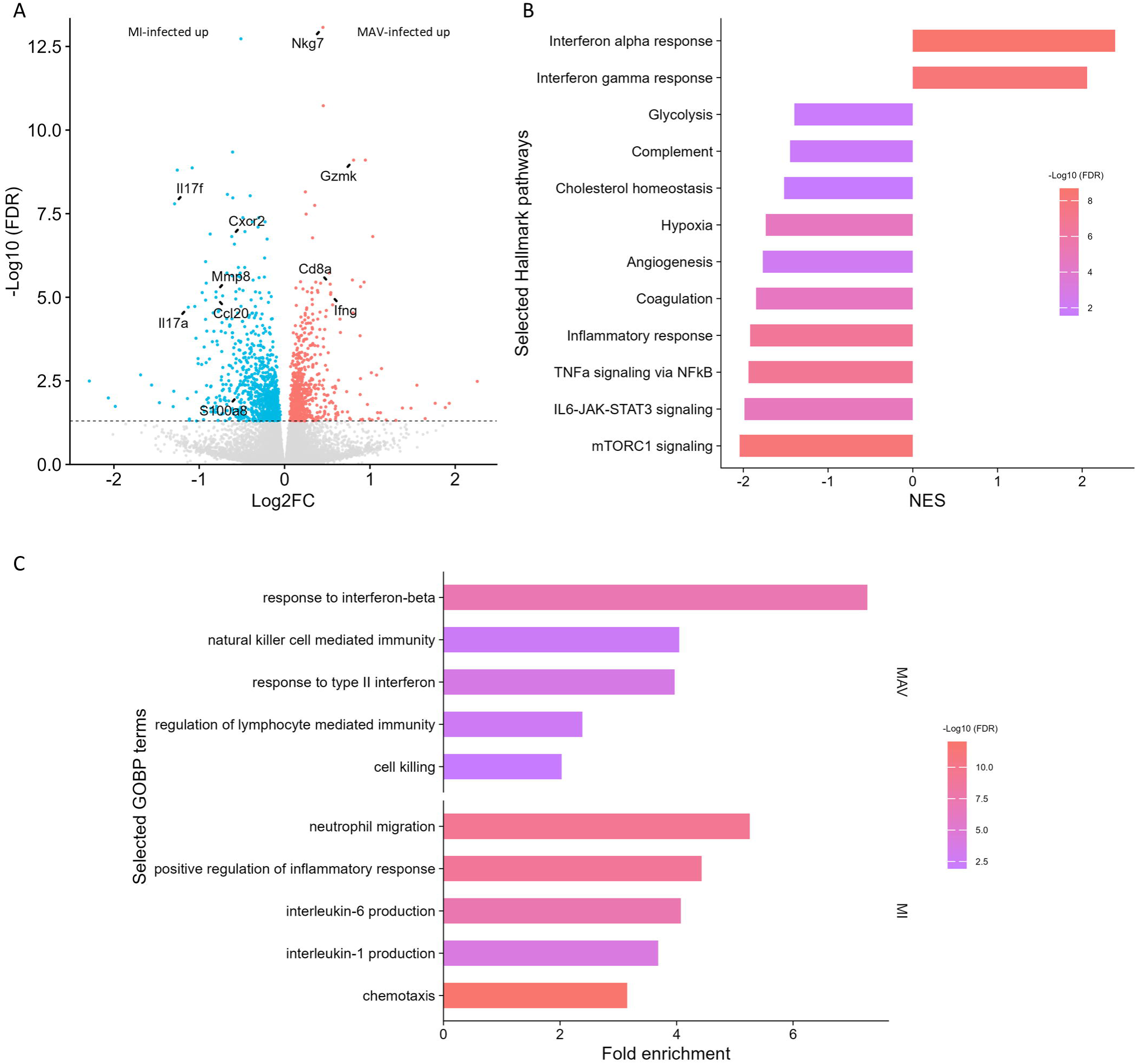
Species-associated transcriptional profiles in lungs infected with MAV and MI. Mice were infected with FKJ-1 (n = 12), FKJ-2 (n = 12), FKJ-3.1 (n = 6), FKJ-5 (n = 12), FKJ-8 (n = 12), FKJ-14 (n = 6), FKJ-27 (n = 6), or NBRC112750 (n = 6) for 8 weeks. Infected lung samples were subjected to bulk RNA-seq. (A) Volcano plot showing differentially expressed genes (DEGs) in MAC-infected lungs. Mouse lungs infected with four MAV isolates or four MI isolates for 8 weeks were subjected to RNA-seq. Red and blue dots indicate upregulated genes in MAV- and MI-infected lungs, respectively. Genes with a false discovery rate (FDR) of less than 0.05 were defined as DEGs. Representative genes associated with IFN-γ/cytotoxicity and IL-17/neutrophilic inflammation are labeled. (B) Gene set enrichment analysis (GSEA) of MAC-infected lungs. Selected Hallmark gene sets associated with immune responses and immunometabolism are depicted. Bar length indicates the normalized enrichment score (NES), with positive and negative NES values representing enrichment in MAV*-* and MI-infected lungs, respectively. Color indicates the −log_10_-transformed FDR. (C) Gene Ontology biological process (GOBP) enrichment analysis of the upregulated DEGs in lungs infected with MAV and MI. Selected immune response-associated GOBP terms are shown for MAV- and MI-infected lungs. Bar length indicates fold enrichment. Color indicates the −log_10_-transformed FDR.

To further characterize these species-associated immune programs, we analyzed MAV- and MI-infected lung transcriptomes using curated gene signatures related to IFN-γ/cytotoxicity and IL-17/neutrophilic inflammation (Fig. 3 and Table S2). Heatmap analysis demonstrated that IL-17/neutrophilic inflammation-associated genes were consistently elevated in MI-infected lungs. In contrast, IFN-γ/cytotoxicity-associated genes exhibited a more heterogeneous pattern, with subsets of both MAV- and MI-infected samples exhibiting high expression levels (Fig. 3A). Signature score analysis at the individual-mouse level revealed higher IFN-γ/cytotoxicity-associated scores in MAV-infected lungs (*P* = 5.03 × 10^−5^) and higher IL-17/neutrophilic inflammation-associated scores in MI-infected lungs (*P* = 3.02 × 10^−5^) (Fig. 3B). When the scores were averaged by isolate, the IL-17/neutrophilic inflammation-associated score remained higher in MI-infected lungs (*P* = 0.049), whereas the IFN-γ/cytotoxicity-associated score did not reach statistical significance (*P* = 0.222) (Fig. 3C). In leave-one-isolate-out analyses, the estimated MI-associated increase in the IL-17/neutrophilic inflammation-associated score remained positive after excluding each isolate in turn, indicating that this association was not driven by a single isolate (Fig. S3). These results suggest that MI infection is associated with a robust IL-17/neutrophilic inflammatory signature. In contrast, IFN-γ/cytotoxicity-associated responses exhibit greater isolate-dependent heterogeneity, despite their relative enrichment in MAV-infected lungs at the individual mouse-level.

**FIG 3.**
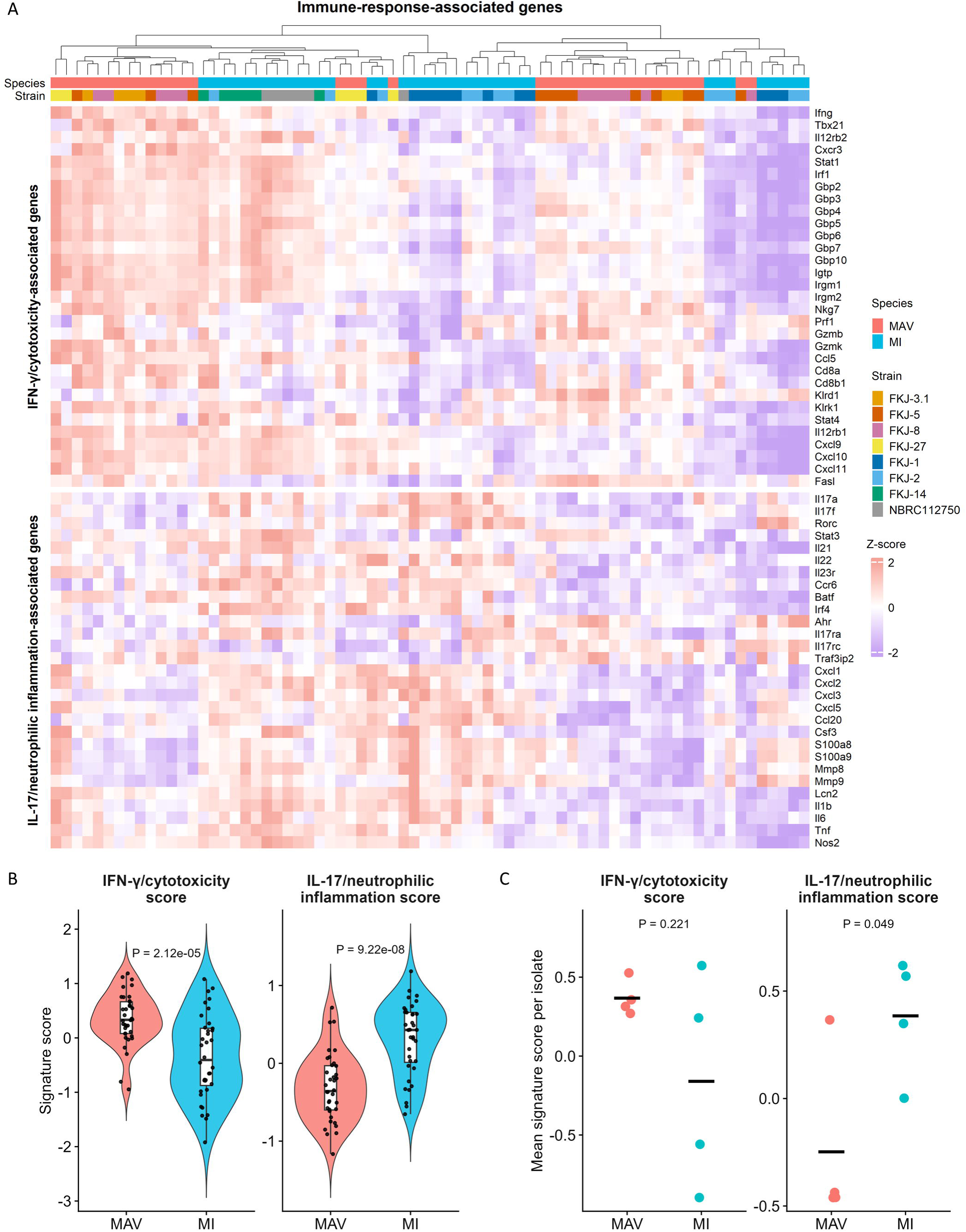
Immune signature profiling of MAC-infected mouse lungs. (A) Heatmap showing the expression profiles of IFN-γ/cytotoxicity-associated genes and IL-17/neutrophilic inflammation-associated genes in MAC-infected mouse lungs. Genes were selected from curated immune response signatures and grouped into IFN-γ/cytotoxicity-associated and IL-17/neutrophilic inflammation-associated modules (Table S2). Columns were clustered across all infected samples without a priori separation by species. (B) Individual mouse-level signature scores for IFN-γ/cytotoxicity-associated genes and IL-17/neutrophilic inflammation-associated genes in MAV- and MI-infected mouse lungs. Each data point represents one mouse. *P* values were determined using Welch’s *t*-test. (C) Isolate-level mean signature scores for IFN-γ/cytotoxicity-associated genes and IL-17/neutrophilic inflammation-associated genes in MAV- and MI-infected mouse lungs. Each data point represents one isolate. *P* values were determined using Welch’s *t*-test.

Because pulmonary growth phenotypes differed among MAC isolates (Figs 1 and S1), we performed a sensitivity analysis including CFU phenotype as a covariate in the edgeR model (Fig. S4 and Table S3). After adjustment for CFU phenotype (Increased: FKJ-1, FKJ-8, FKJ-27, and NBRC112750. Maintained: FKJ-2, FKJ-3.1, FKJ-5, and FKJ-14), the MI-associated IL-17/neutrophilic program remained evident, including *Il17a*, *Il17f*, *Cxcl2*, *Cxcl3*, *Cxcl5*, *Cxcr2*, *S100a8*, *S100a9*, *Mmp8*, and *Mmp9*. In contrast, MAV-associated differences were retained mainly in cytotoxic/NK-cell-related genes, including *Nkg7*, *Gzmb*, *Gzmk*, *Prf1*, *Fasl*, and *Cd8a/b*, whereas canonical interferon-stimulated genes were less consistently retained after adjustment. These results suggest that the MI-associated IL-17/neutrophilic signature is not explained solely by pulmonary growth phenotype.

### Validation of immune signatures and cellular composition using representative isolate pair FKJ-1 and FKJ-8

To validate the RNA-seq–based immune signatures in an independent cohort, we examined the expression of genes associated with IFN-γ/cytotoxic responses and IL-17/neutrophilic inflammation in the lungs of mice infected with FKJ-1 or FKJ-8 using RT-qPCR (Fig. 4A, B). FKJ-1 and FKJ-8 were selected as the representative MI–MAV isolate pair for detailed analysis based on two criteria. First, FKJ-1 and FKJ-8 exhibited similar proliferative kinetics in the murine pulmonary infection model at 8 weeks p.i. (Fig. S1), as described previously (19, 20). Second, the comparison of FKJ-1- and FKJ-8-infected lungs reproduced the overarching transcriptional trends observed in the broader comparison between MAV- and MI-infected lungs (Fig. S2C). Thus, FKJ-1 and FKJ-8 provided a suitable isolate pair for evaluating these species-associated immune signatures under comparable levels of pulmonary bacterial persistence. The specific genes for validation were selected from the DEGs associated with IFN-γ/cytotoxic responses or IL-17/neutrophilic inflammation in MAC-infected lungs. RT-qPCR analysis confirmed that their expression patterns in FKJ-1- and FKJ-8-infected lungs were consistent with the bulk RNA-seq results.

**FIG 4.**
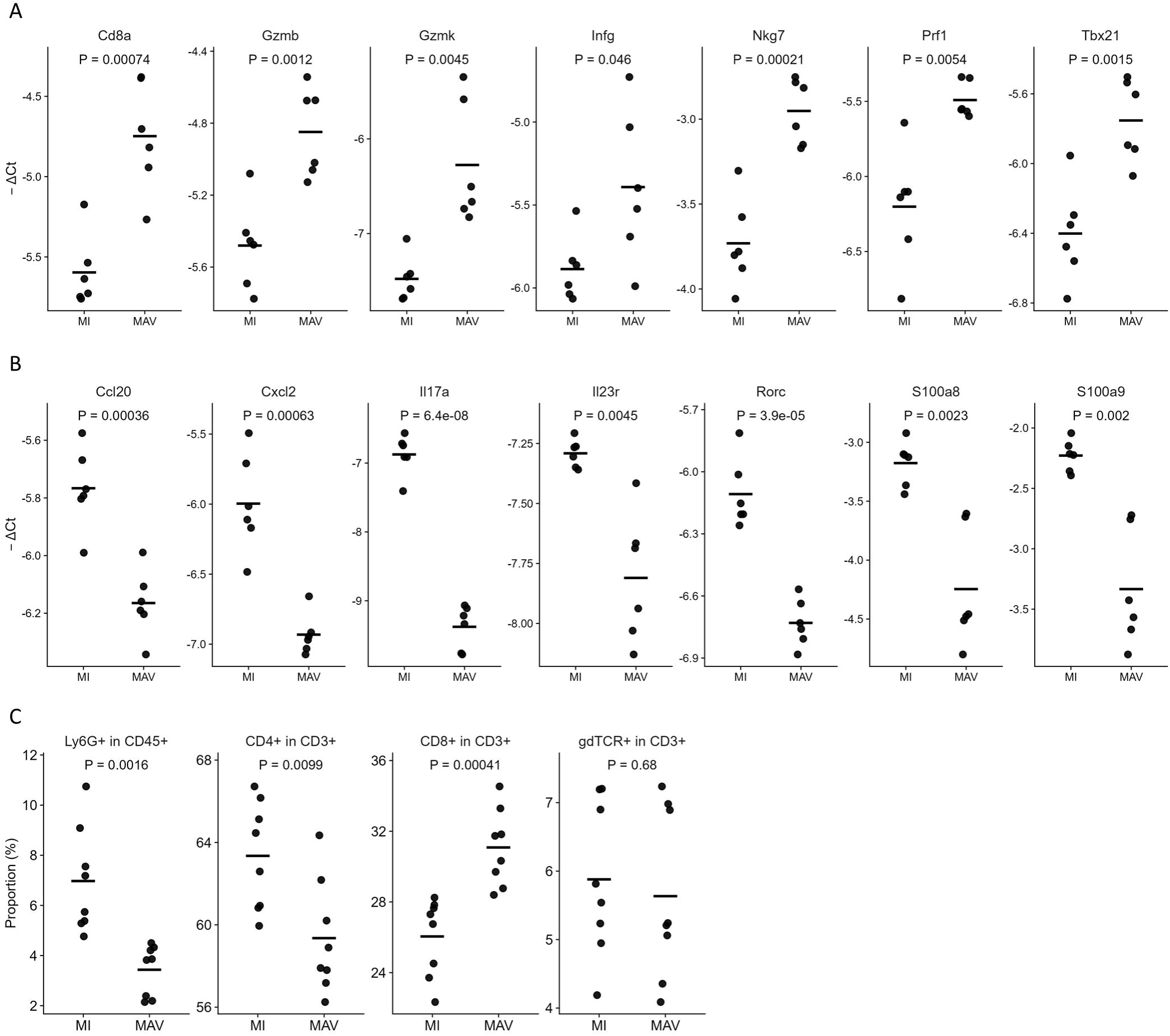
Validation of gene expression and cellular composition in MAC-infected mouse lungs. (A, B) RT-qPCR analysis of IFN-γ/cytotoxic-associated genes (A) and IL-17/neutrophilic inflammation-associated genes (B) in MAC-infected mouse lungs. Mouse lungs infected with FKJ-1 (MI, n = 6) or FKJ-8 (MAV, n = 6) for 8 weeks were subjected to RT-qPCR. The genes associated with these immune signatures were selected from DEGs identified *via* RNA-seq. −ΔCT values were calculated as −(CT value of the indicated gene − CT value of *Rplp1*). (C) Flow cytometric analysis of MAC-infected lungs. Mouse lungs infected with FKJ-1 (MI; n = 8) or FKJ-8 (MAV; n = 8) for 8 weeks were analyzed for the indicated immune cell populations. Each data point represents one mouse. *P* values were determined using Welch’s *t*-test.

Flow cytometric analysis was subsequently performed to examine the cellular composition of lungs infected with FKJ-1 and FKJ-8 (Fig. 4C). The proportion of neutrophils was higher in FKJ-1-infected lungs than in FKJ-8-infected lungs, consistent with the enrichment of neutrophil recruitment-related transcriptional programs in MI-infected lungs. Among the T-cell populations, CD4^+^ T cells were more abundant in FKJ-1-infected lungs, whereas CD8^+^ T cells were more abundant in FKJ-8-infected lungs. In contrast, the proportions of γδTCR^+^ T cells were comparable between the two groups. Moreover, transcriptomic analysis also revealed the upregulation of *Ccl20* and *Ccl17/Ccl22*, chemokine genes associated with CCR6^+^ Th17-type cells and CCR4^+^ CD4^+^ T-cells (25, 26), respectively, in FKJ-1- and other MI-infected lungs (Fig. S5). Collectively, these cellular profiles were broadly consistent with the transcriptomic findings that FKJ-1 infection is associated with IL-17/neutrophilic inflammation, whereas FKJ-8 infection is characterized by IFN-γ/cytotoxic immune responses.

### Single-cell landscape of lungs infected with FKJ-1 and FKJ-8

We applied scRNA-seq to determine the cellular basis of the distinct immune transcriptional programs observed between FKJ-1- and FKJ-8-infected lungs. BALB/c mice were infected for 8 weeks with FKJ-1 (MI) or FKJ-8 (MAV), the representative isolate pair selected for detailed analysis. Lung cells were then isolated and subjected to scRNA-seq. From the 125,950 cells isolated from FKJ-1- or FKJ-8-infected lung tissue, 12 major cell types were manually annotated (Fig. 5A, B, and Table S4). Diverse immune cell populations were identified, which included T cells, natural killer (NK) cells, macrophages, neutrophils, conventional dendritic cells (cDCs), plasmacytoid dendritic cells (pDCs), basophils, B cells, and plasma cells. The overall immune cell composition was broadly similar between FKJ-1- and FKJ-8-infected lungs (Fig. 5C); however, neutrophils and NK cells were more abundant in FKJ-1- and FKJ-8-infected lungs, respectively (Fig. 5D). These results were consistent with the RT-qPCR and flow cytometric analyses (Fig. 4).

**FIG 5.**
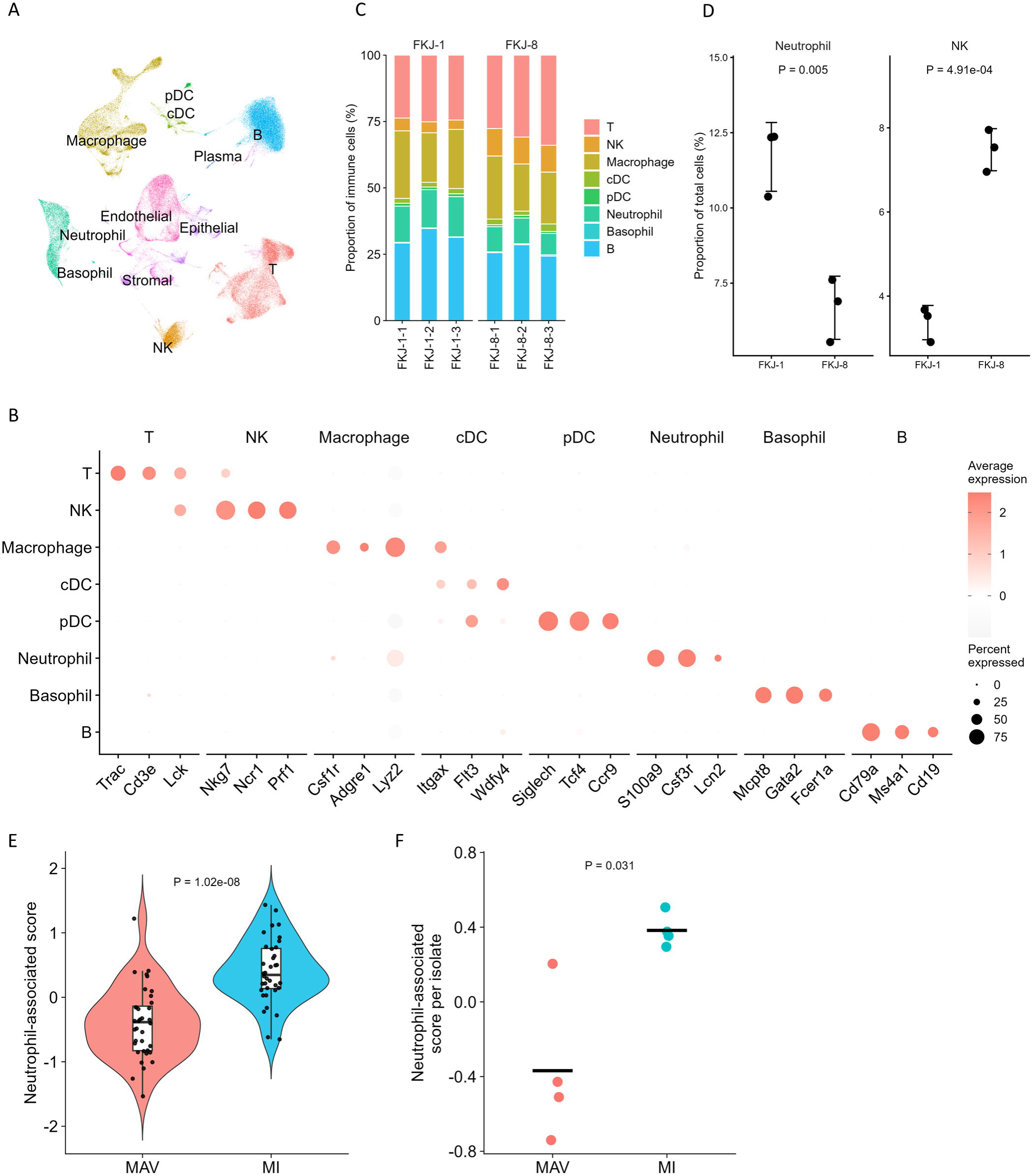
Single-cell transcriptomic landscape of MAC-infected mouse lungs. (A) Uniform manifold approximation and projection (UMAP) plot of 125,950 cells isolated from FKJ-1-infected mouse lungs (MI; n = 3) or FKJ-8 (MAV; n = 3). In total, 12 major cell types are shown: T cells, natural killer (NK) cells, macrophages, conventional dendritic cells (cDCs), plasmacytoid dendritic cells (pDCs), neutrophils, basophils, B cells, plasma cells, epithelial cells, endothelial cells, and stromal cells. (B) Dot plot showing the expression of selected marker genes across the major immune cell populations. Dot size indicates the percentage of cells expressing each gene. Color indicates the average expression level. (C) Stacked bar plot showing the proportions of immune cell populations in lungs infected with FKJ-1 or FKJ-8. Each bar represents one mouse. Colors indicate the respective immune cell populations. (D) Proportions of neutrophils and NK cells in lungs infected with FKJ-1 or FKJ-8. The immune cell populations that differed significantly between FKJ-1- and FKJ-8-infected lungs are shown. Each dot represents one mouse. *P* values were determined using Welch’s *t*-test. (E, F) Neutrophil-associated scores among the MAC isolates. Neutrophil-associated scores were calculated from bulk RNA-seq data using marker genes derived from the scRNA-seq neutrophil cluster. Scores are shown at the individual-mouse level (E) and isolate level (F). Each data point represents one mouse (E) or one isolate (F). In (E), violin plots show the data distribution, and box plots indicate the median and interquartile range. In (F), horizontal bars indicate the mean. *P* values were determined using Welch’s *t*-test.

To determine whether the neutrophil program identified using scRNA-seq was reflected across the broader bulk RNA-seq cohort, we projected a neutrophil-associated gene signature derived from the scRNA-seq neutrophil cluster onto the bulk RNA-seq profiles from eight MAC isolates. MI-infected lungs exhibited higher neutrophil-associated transcriptional scores than MAV-infected lungs at both the individual-mouse and isolate levels (Fig. 5E, F), supporting a broader association of MI infection with neutrophilic pulmonary inflammation.

### T cell and macrophage states in FKJ-1- and FKJ-8-infected lungs

We next analyzed T-cell subclusters in MAC-infected lungs (Fig. 6A, B, and Table S5). Among CD4^+^ T cells, Th1, Th17, and regulatory T (Treg) cell clusters were identified, consistent with previous reports describing the pulmonary accumulation of CD4^+^ Th1, Th17, and Treg cells during MAC infection (11, 13). We also identified an *Il17a*-expressing γδ T-cell cluster (γδTh17) in MAC-infected lungs. This cluster expressed *Pdcd1*, a phenotype similarly observed in the necrotizing granulomas of *M. tuberculosis*-infected C3HeB/FeJ mice (27). Analysis of T-cell cluster abundance demonstrated that CD4^+^ Th17 cells were significantly increased in FKJ-1-infected lungs, whereas γδTh17 cell abundance did not significantly differ between FKJ-1- and FKJ-8-infected lungs (Fig. 6C). In contrast, intracellular cytokine staining revealed that the proportions of IL-17A-expressing cells among both CD4^+^ T cells and γδTCR^+^ T cells were significantly higher in FKJ-1-infected lungs than in FKJ-8-infected lungs (Fig. 6D). These phenotypic findings were consistent with the bulk RNA-seq results, showing enrichment of IL-17/neutrophilic inflammatory signatures in MI-infected lungs (Figs. 2 and 3).

**FIG 6.**
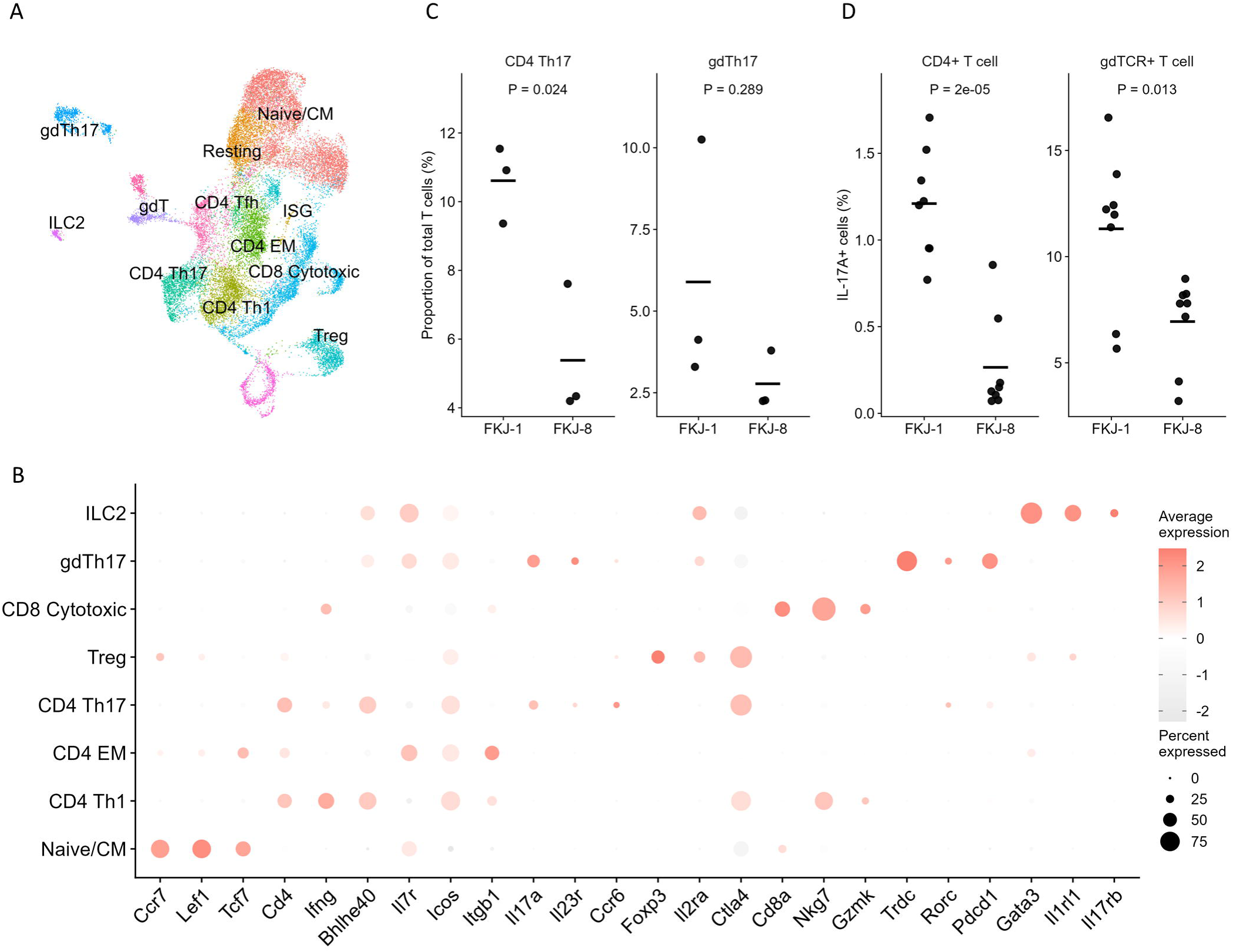
T-cell clusters in MAC-infected mouse lungs. (A) UMAP plot showing T-cell populations in MAC-infected mouse lungs. T cells and other lymphoid cells were subsetted from total MAC-infected lung cells and reclustered. The following populations are shown: naïve/central memory (CM) T cells, resting T cells, interferon-stimulated gene (ISG) T cells, CD4^+^ Th1 cells, CD4^+^ effector memory (EM) T cells, CD4^+^ Th17 cells, CD4^+^ Tfh cells, regulatory T cells (Treg), CD8^+^ cytotoxic T cells, γδ T cells, γδ Th17 cells, and ILC2s. Cycling T cells and contaminating clusters are excluded. (B) Dot plot showing the expression of selected marker genes across the major T-cell populations. Dot size indicates the percentage of cells expressing each gene. Color indicates the average expression level. (C) Proportions of CD4^+^ Th17 and γδ Th17 cells within the T-cell compartment. The proportions of CD4^+^ Th17 and γδ Th17 cells among total T cells are shown for FKJ-1- and FKJ-8-infected lungs. (D) Validation of IL-17A-expressing cells among CD4^+^ T cells and γδ T cells. Intracellular cytokine staining for IL-17A was performed using lung cells from mice infected with FKJ-1 (n = 8) or FKJ-8 (n = 8) for 8 weeks. The proportions of IL-17A-positive cells among CD4^+^ T cells or γδ T cells are shown. Each data point represents one mouse. *P* values were determined using Welch’s *t*-test.

A detailed cluster analysis of the macrophage and monocyte populations in MAC-infected lungs was performed (Fig. 7A, B, and Table S6). These populations were classified into classical monocytes, patrolling monocytes, alveolar macrophages (AMs), interstitial macrophages (IMs), and infection-associated macrophage populations based on the expression of established marker genes. These included *Ly6c2* and *Ccr2* for classical monocytes, *Nr4a1* and *Ace* for patrolling monocytes, *Siglecf* for AMs, and *C1q* for IMs. *Spp1* and *Trem2* were used as additional markers to identify infection-associated or inflammatory interstitial-like macrophage states (28–32).

**FIG 7.**
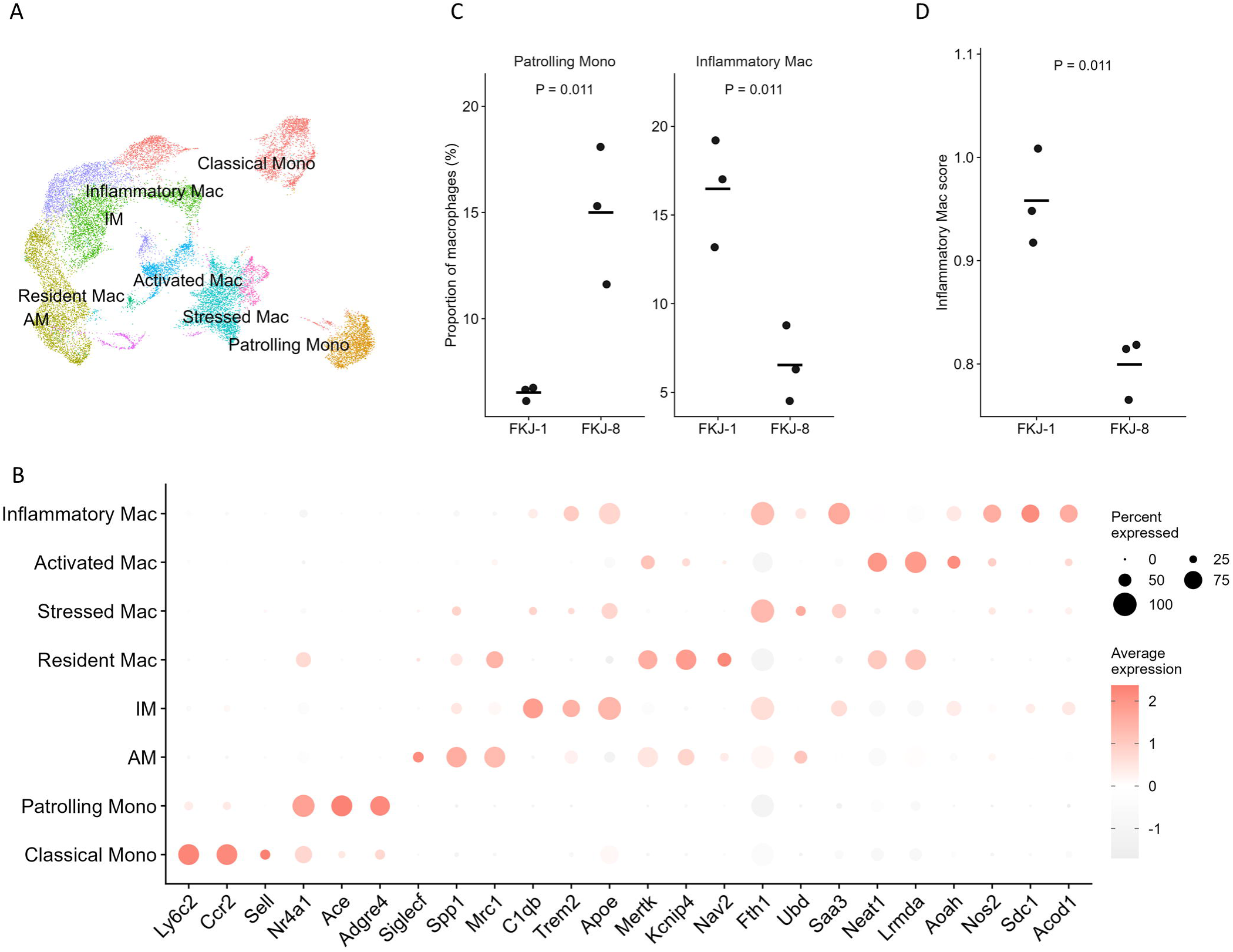
Macrophage clusters in MAC-infected mouse lungs. (A) UMAP plot showing macrophage and monocyte populations in MAC-infected mouse lungs. Macrophages and monocytes were subsetted from the total MAC-infected lung cells and reclustered. The following populations are shown: classical monocytes (Mono), patrolling monocytes, alveolar macrophages (AM), interstitial macrophages (IM), resident-like macrophages (Resident Mac), stressed macrophages, activated macrophages, and inflammatory macrophages. Cycling and contaminating clusters are excluded. (B) Dot plot showing the expression of selected marker genes across the macrophage and monocyte populations. Dot size indicates the percentage of cells expressing each gene. Color indicates the average expression level. (C) Proportions of patrolling monocytes and inflammatory macrophages within the macrophage/monocyte compartment. The proportions of patrolling monocytes and inflammatory macrophages among total macrophages/monocytes are shown for FKJ-1- and FKJ-8-infected lungs. (E) Inflammatory macrophage-associated gene scores in the inflammatory macrophage cluster. Scores were calculated using the marker genes of inflammatory macrophages. Each data point represents one mouse. Horizontal bars indicate the mean. *P* values were determined using Welch’s *t*-test.

Macrophage populations were further annotated into resident-like, stressed, activated, and inflammatory macrophages based on cluster-enriched marker genes (Fig. 7A, B). Resident-like, stressed, and activated macrophage clusters were characterized by the relatively high expression of *Mertk* and *Kcnip4*, *Fth1* and *Ubd*, and *Neat1* and *Lrmda*, respectively (33–37). Inflammatory macrophages were distinguished by the prominent expression of *Nos2* and additional inflammation-associated genes, including *Saa3*, *Sdc1*, and *Acod1*. These precise annotations identified distinct macrophage activation states within MAC-infected lungs.

Finally, we compared the proportions of macrophage and monocyte clusters between FKJ-1- and FKJ-8-infected lungs, finding that inflammatory macrophages and patrolling monocytes were significantly more abundant in FKJ-1- and FKJ-8-infected lungs, respectively (Fig. 7C). Further, we assessed inflammatory macrophage-associated gene signature scores within the inflammatory macrophage cluster (Fig. 7D). The expression score, calculated from the top 10 DEGs of inflammatory macrophages (Table S2), was significantly higher in FKJ-1-infected than in FKJ-8-infected lungs. Together, these results suggest that, compared to FKJ-8 infection, FKJ-1 infection is associated with an increased abundance of inflammatory macrophages and an enhanced inflammatory gene expression signature within those macrophages.

## Discussion

In this study, we established and characterized a panel of MAC clinical isolates capable of establishing persistent pulmonary infection in BALB/c mice. Using this model, we compared the pulmonary host immune responses induced by MAV and MI. Compared to uninfected control lungs, both MAV and MI infections induced common inflammatory and IFN-related transcriptional responses; however, a direct comparison between MAV- and MI-infected lungs revealed distinct, species-associated immune programs. MI-infected lungs exhibited robust enrichment of IL-17/neutrophilic inflammatory signatures. In contrast, IFN-γ/cytotoxicity-associated responses were relatively enriched in MAV-infected lungs but displayed greater isolate-dependent heterogeneity. These transcriptional differences were supported by RT-qPCR, flow cytometry, and scRNA-seq analyses. Together, these findings suggest that among MAC clinical isolates capable of persistent pulmonary infection in BALB/c mice, MAV and MI natively elicit different balances of pulmonary immune responses.

Although patients with MAC-PD display clinical heterogeneity (2, 38, 39), it remains unclear whether this heterogeneity is driven by differences in the magnitude or balance of pulmonary immune programs induced by different MAC species or strains. Previous murine studies have demonstrated that MAC isolates inherently differ in their capacity to persist, proliferate, and induce lung pathology (19–22), suggesting that bacterial factors influence disease progression. To address whether such bacterial differences are reflected in host immune responses, we compared the pulmonary transcriptomes of mice infected with MAV and MI isolates under controlled experimental conditions (Fig. 2). Importantly, MAV and MI did not induce mutually exclusive immune responses; rather, they activated broadly shared inflammatory programs but differed in the relative dominance of specific immune modules. A sensitivity analysis further indicated that the MI-associated IL-17/neutrophilic program was retained at the isolate level and after adjustment for pulmonary growth phenotype (Fig. 3 and Fig. S4), supporting the interpretation that this signature was not explained solely by increased bacterial proliferation. In contrast, broad interferon-related transcriptional programs were less consistently retained after isolate-level analysis and adjustment for pulmonary growth phenotype, suggesting that these responses may reflect both species-associated differences and the inflammatory burden imposed by proliferative isolates. Together, these findings suggest that species- or strain-associated differences in MAC-PD may manifest not only through bacterial persistence or lesion development but also through the balance of pulmonary immune responses.

Th1 and Th17 responses have been widely implicated in the pathogenesis of MAC infection, and the balance between these immune axes may influence disease progression (10–13). In this study, MI-infected lungs exhibited stronger IL-17/neutrophilic inflammatory signatures than MAV-infected lungs (Fig. 2 and Fig. 3). This transcriptional bias was functionally reflected at the cellular level, as FKJ-1-infected lungs contained higher proportions of neutrophils than FKJ-8-infected lungs (Fig. 4). Moreover, the projection of a neutrophil-associated gene signature, derived from the scRNA-seq neutrophil cluster, onto the bulk RNA-seq cohort revealed higher neutrophil-associated transcriptional scores in MI-infected lungs across multiple isolates (Fig. 5). Collectively, these findings suggest that, in this murine MAC-PD model, MI possesses a greater tendency than MAV to induce neutrophil-associated pulmonary inflammation.

Neutrophil-associated inflammation has been clinically and experimentally linked to disease severity in NTM-PD, including MAC-PD (40). A whole-blood transcriptomic study of patients with MAC-PD previously identified a severe clinical cluster characterized by neutrophil activation, suggesting that systemic neutrophil-associated signatures may serve as biomarkers for clinically severe disease states (15). In murine models, virulent MI strains have similarly been associated with higher pulmonary bacterial burdens and increased neutrophil infiltration (21). IL-17A-mediated host responses, including those involving IL-17A-producing CD4^+^ T cells and γδ T cells, have been implicated in protective immunity against pulmonary mycobacterial infection (41, 42). However, the role of this pathway appears to be context-dependent. In a chronic MI infection model, for instance, neutrophil depletion did not substantially improve disease outcomes, whereas early neutralization of IL-17A reduced chronic bacterial burdens, lung damage, and mortality (10). These findings suggest that, in chronic MI-associated pulmonary infections, IL-17A-driven inflammatory programs may exacerbate disease progression, at least in part, through mechanisms linked to neutrophilic inflammation rather than through neutrophil accumulation alone. In our model, although FKJ-1 and FKJ-8 yielded comparable pulmonary bacterial burdens at 8 weeks p.i. (Fig. S1), FKJ-1-infected lungs exhibited stronger IL-17/neutrophilic inflammatory signatures and higher neutrophil proportions (Fig. 4). Moreover, neutrophil-associated transcriptional scores projected onto the bulk RNA-seq cohort were higher in MI-infected lungs across multiple isolates (Fig. 5). These observations suggest that MI infection possesses a greater tendency than MAV infection to induce IL-17–neutrophil-skewed pulmonary inflammatory environment in this murine MAC-PD model, even when bacterial persistence remains comparable between representative strains. Given that MAC can induce neutrophil extracellular trap formation and MMP-8/MMP-9 release from neutrophils (43), neutrophil-associated inflammation may be directly associated with tissue remodeling and destructive lesion formation in MAC-PD (20, 44, 45). However, whether IL-17–neutrophil axis-associated programs promote the formation of necrotizing granuloma or cavitary disease in MAC-PD requires further investigation.

In addition to lymphoid and neutrophilic responses, scRNA-seq analysis revealed an expansion of *Nos2*-expressing inflammatory macrophages and higher inflammatory macrophage-associated gene scores in FKJ-1-infected lungs (Fig. 7). Because NOS2-dependent nitric oxide is generally associated with antimicrobial macrophage activation during mycobacterial infection (46), the accumulation of *Nos2*^+^ inflammatory macrophages may be associated with a host response attempting to control persistent infection. However, in MAC infections, NOS2-derived nitric oxide regulates granuloma formation without reducing bacterial burdens (47). Consistent with this complexity, the expansion of *Nos2*^+^ inflammatory macrophages should not be interpreted simply as evidence of effective bacterial control. Instead, this population may reflect a complex inflammatory myeloid state in which antimicrobial and tissue-remodeling programs are strongly induced during persistent infection.

The inflammatory macrophage states identified in our MAC model also shared certain features with the proinflammatory macrophage states described in *M. tuberculosis*-infected lungs. Pisu et al. reported that *Nos2* upregulation in infected macrophages was associated with stressed *M. tuberculosis* and that these proinflammatory macrophage subsets expressed a distinct set of genes, including *Nos2*, *Clec4e*, *Saa3*, *Ccl5*, and *Ubd* (48). In our MAC model, MAC infection was similarly associated with the expansion of *Nos2*^+^ inflammatory macrophages expressing overlapping inflammation-associated genes, including *Clec4e*, *Saa3*, and *Ccl5*. These similarities suggest that *Nos2*^+^ inflammatory macrophage states may represent a conserved inflammatory macrophage program induced across mycobacterial infection, although their functional consequences may differ between *M. tuberculosis* and MAC infections. Further studies are required to determine whether these macrophages contribute primarily to bacterial control, tissue injury, granulomatous remodeling, or a combination of these processes.

A notable limitation of this study is that the MAV and MI isolates analyzed here were not randomly selected to represent the full clinical diversity of each species. Rather, the clinical MAC isolates used in this study were selected based on their ability to maintain or increase pulmonary bacterial burdens in BALB/c mice. Therefore, the species-associated transcriptional differences identified herein must be interpreted in the context of this selected panel of persistent MAC clinical isolates. In addition, the RT-qPCR, flow cytometric, and scRNA-seq analyses were performed using FKJ-1 and FKJ-8 as the representative isolate pair, selected on the basis of their comparable bacterial persistence and concordant bulk transcriptional trends. Thus, the distinct cellular phenotypes identified in FKJ-1- and FKJ-8-infected lungs likely reflect an interplay of both species-associated and isolate-specific effects.

In conclusion, this study demonstrates that the MAV and MI isolates included in the eight-isolate MAC panel induced broadly shared, yet quantitatively distinct, pulmonary immune programs in a chronic murine MAC-PD model. MI-infected lungs exhibited enhanced IL-17/neutrophilic inflammatory signatures, whereas IFN-γ/cytotoxicity-associated responses were relatively enriched in MAV-infected lungs, albeit with observable isolate-dependent heterogeneity. Deeper analysis using a representative isolate pair further corroborated that, compared with FKJ-8 (MAV) infection, FKJ-1 (MI) infection was associated with increased proportions of IL-17A-producing lymphocytes and an expansion of *Nos2*^+^ inflammatory macrophage states. These findings strongly suggest that species-associated differences in the host immune responses may contribute to the clinical and pathological heterogeneity of MAC-PD. The well-characterized panel of persistent MAC clinical isolates established in this study serves as a useful preclinical platform for dissecting host–pathogen interactions and evaluating future therapeutic strategies in chronic pulmonary MAC infections.

## MATERIALS AND METHODS

### Ethics statement

The animal experiments conducted in this study were approved by the Animal Care and Use Committee of the Research Institute of Tuberculosis (RIT) (Permit number: No. 2023-02). All animals were handled in strict accordance with the ethical guidelines of the RIT. The endpoints were set to determine whether mice required euthanasia due to severe distress or impending mortality from MAC infection, defined primarily by a bodyweight loss of >20% of their initial weight at the time of infection.

### Mice

Female BALB/c mice (6 weeks old) were purchased from Japan SLC, and transferred to the biosafety level II animal facility of the RIT. The mice were maintained in negative-pressure ventilated racks and provided with sterile bedding, water, and standard chow ad libitum throughout the experimental period.

### Infection with MAC isolates

MAC clinical isolates, obtained from the sputum of patients with MAC-PD visiting Fukujuji Hospital (Japan Anti-Tuberculosis Association, Tokyo, Japan), were cultured at 37°C in 7H9 medium supplemented with 10% Middlebrook ADC (BD Biosciences), 0.5% casamino acids, and 0.05% Tween 80 (mycobacterial medium) (49). All patients met the diagnostic criteria set forth by the 2020 guidelines for NTM-PD treatment (1). The established MAC isolates FKJ-1, FKJ-2, FKJ-5, FKJ-8, and NBRC112750 (19, 20) were also cultured at 37°C in mycobacterial medium. The screening of clinical MAC isolates exhibiting persistent phenotypes in mouse lungs was performed as described previously (19). Single-cell suspensions were prepared as described previously (19) and stored at −80°C until use. For infection experiments, the frozen stocks were thawed and maintained on ice until inoculation, without prior broth subculture. Mice were infected with the MAC strains intranasally at a dose of 2 × 10^6^ CFU in 60 μL of sterile water.

### Genome sequencing and phylogenetics

Genomic DNA from the MAC isolates and the corresponding sequence libraries were prepared as described previously (19). Phylogenetic trees were constructed using the maximum likelihood method implemented in REALPHY software version 1.1.3 with default parameters (50). The GenBank genome sequences of *M*. *avium* 104 (accession number: CP000479) and *M*. *intracellulare* ATCC 13950 (accession number: CP003322) were used as reference genomes. WGS data for the MAC isolates FKJ-1, FKJ-2, FKJ-5, FKJ-8, and NBRC112750 were accessed from the DDBJ database (accession numbers: PRJDB12771 and PRJDB37637) (19, 20).

### Collection of lung lobes and bulk RNA-seq

At 8 weeks p.i., the infected mice were anesthetized *via* intraperitoneal administration of an anesthetic cocktail containing 0.75 mg/kg medetomidine, 4 mg/kg midazolam, and 5 mg/kg butorphanol. The mice were euthanized by exsanguination, and their lungs were excised. For CFU assays, whole lungs were homogenized, serially diluted, and plated on 7H10 or 7H11 agar plates supplemented with 10% Middlebrook OADC enrichment (BD Biosciences) and 0.5% glycerol.

For bulk RNA-seq, mice were infected with FKJ-1 (n = 12), FKJ-2 (n = 12), FKJ-3.1 (n = 6), FKJ-5 (n = 12), FKJ-8 (n = 12), FKJ-14 (n = 6), FKJ-27 (n = 6), or NBRC112750 (n = 6) for 8 weeks. Uninfected control mice were processed in parallel (n = 12). Whole lung lobes were stored in RNA preservation solution (51) at −80°C until extraction. RNA isolation from the lung lobes and subsequent bulk RNA-seq were performed as described previously (52). Libraries were sequenced on an Illumina NextSeq 1000 platform to generate approximately 20 million 2 × 50-bp or 2 × 100-bp paired-end reads per library.

### scRNA-seq

For scRNA-seq, mice infected with FKJ-1 (n = 3) or FKJ-8 (n = 3) for 8 weeks were euthanized by exsanguination, and their lungs were excised after cardiac perfusion with PBS. Single-cell suspensions were prepared from the lung tissue using a Lung Dissociation Kit (Miltenyi Biotec). scRNA-seq libraries were constructed using the Illumina Single Cell 3′ RNA Prep T10 kit (Illumina), and the resulting libraries were sequenced on an Illumina NextSeq 1000 platform according to the manufacturer’s instructions.

### Bioinformatics

For bulk RNA-seq, the raw sequencing data were processed as described previously (53), with slight modifications. Briefly, raw reads were quality-trimmed using Trim Galore (version 0.6.10) (https://github.com/FelixKrueger/TrimGalore). The processed reads were then aligned to the mouse reference genome (mm39) using STAR (version 2.7.11b) (54) and gene-level counts were quantified using featureCounts (version 2.1.1) (55). Differential gene expression analysis was performed using edgeR (version 4.8.2) (56) based on generalized linear models and quasi-likelihood F-tests (57). GSEA (58) was performed using fgsea (version 1.36.2) (59) leveraging Hallmark gene sets obtained from the Molecular Signatures Database (MSigDB). Functional characterization of DEGs was achieved by GOBP enrichment analysis using clusterProfiler (version 4.18.4) (60) and ShinyGO (version 0.85.2) (61).

For scRNA-seq, the raw sequencing reads were aligned to the mouse reference genome (mm39) using PIPseeker version 3.3 (Fluent Biosciences). Subsequent analyses were performed in R (version 4.5) using the Seurat package (version 5.4) (62). For each sample, low-quality cells were filtered out based on the following criteria: cells with fewer than 300 or more than 4,000 detected genes, fewer than 400 UMIs, or more than 15% mitochondrial gene content were excluded. Doublets were identified for each sample using scDblFinder, and only cells classified as singlets were retained for downstream analysis. After quality control and doublet removal, all samples were merged into a single Seurat object. The RNA assay was normalized using the NormalizeData function, variable features were identified using FindVariableFeatures, and the data were scaled using ScaleData. Principal component analysis was then performed using the first 50 principal components. To correct for sample-dependent effects, the RNA assay was split by sample and integrated using reciprocal principal component analysis (RPCA) integration *via* the Seurat v5 IntegrateLayers function. The integrated RPCA reduction was used for downstream dimensionality reduction and clustering. Shared nearest-neighbor graphs were constructed using the first 30 integrated dimensions, clusters were identified with a resolution of 0.6, and Uniform manifold approximation and projection visualization was performed using the corresponding 30 dimensions.

Signature scores were calculated as the mean *z*-score of the signature genes using log_2_(TPM + 1) values for bulk RNA-seq data, or normalized expression values for scRNA-seq data. Genes not detected in the expression matrix were excluded. Genes used for the estimation of signature scores are listed in Table S2. For comparisons across MAC isolates, signature scores were averaged within each isolate and then compared between MAV and MI isolates using Welch’s *t*-test at both the individual-mouse and isolate levels. Leave-one-isolate-out analyses were performed by excluding one isolate at a time and recalculating the difference in isolate-level mean signature scores between MI and MAV isolates.

### RT-qPCR and flow cytometric analysis

For independent validation experiments, mice were infected with FKJ-1 or FKJ-8 for 8 weeks and subjected to RT-qPCR or flow cytometric analyses. For RT-qPCR analysis, total RNA was isolated from mouse lungs infected with FKJ-1 (n = 6) or FKJ-8 (n = 6), followed by RT-qPCR using the primer sets listed in Table S7. For flow cytometric analysis, lung cell suspensions (FKJ-1; n = 8, FKJ-8; n = 8) were prepared using a Lung Dissociation Kit (Miltenyi Biotec) according to the manufacturer’s instructions. Cells were incubated with an Fc receptor blocking antibody (BioLegend) and subsequently stained with the antibodies listed in Table S8. Stained cells were fixed with Fixation Buffer (BioLegend) for at least 12 h at 4°C prior to acquisition. Cells were analyzed using a BD FACSLyric flow cytometer.

## Data availability

The sequencing data have been deposited in DDBJ under BioProject accession numbers PRJDB42275, PRJDB42278, and PRJDB42283, and in the DDBJ Genomic Expression Archive under accession number E-GEAD-1252.

## ACKNOWLEDGMENTS

This study was supported by the Emerging/Re-emerging Infectious Diseases Project of the Japan Agency for Medical Research and Development (JP23gm1610013, JP23fk0108673, JP23fk0108674, JP23fk0108703, JP25fk0108730, and JP26fk0108750).

We thank Ms. Miyako Seto, Mr. Masanari Matsuda, Mr. Akihiro Hata, Mr. Yoshitomo Baba of the Department of Pathophysiology and Host Defense, RIT, and Ms. Kazue Mizuno of Fukujuji Hospital for their technical support.

## Notes

### Competing Interest Statement

The authors have declared no competing interest.

